# Simu-dependent clearance of dying cells regulates macrophage function and inflammation resolution

**DOI:** 10.1101/312496

**Authors:** Hannah Grace Roddie, Emma Louise Armitage, Simon Andrew Johnston, Iwan Robert Evans

**Affiliations:** Department of Infection, Immunity and Cardiovascular Disease, University of Sheffield, UK; The Bateson Centre, University of Sheffield, UK

## Abstract

Macrophages encounter and clear apoptotic cells during normal development and homeostasis, including at numerous sites of pathology. Clearance of apoptotic cells has been intensively studied, but the effects of macrophage-apoptotic cell interactions on macrophage behaviour are poorly understood. Using *Drosophila* embryos, we have exploited the ease of manipulating cell death and apoptotic cell clearance in this model to identify that the loss of the apoptotic cell clearance receptor Simu leads to perturbation of macrophage migration and inflammatory responses via pathological levels of apoptotic cells. Removal of apoptosis ameliorates these phenotypes, while acute induction of apoptosis phenocopies these defects and reveals that phagocytosis of apoptotic cells is not necessary for their anti-inflammatory action. Furthermore, Simu is necessary for clearance of necrotic debris and rentention of macrophages at wounds. Thus, Simu is a general detector of damaged self and represents a novel molecular player in controlling resolution of inflammation.

---

During development and throughout life, cells are eliminated by programmed cell death and rapidly cleared by phagocytes such as macrophages and glia. Failures in apoptotic cell clearance (efferocytosis) are thought to contribute to disease progression in multiple chronic inflammatory conditions (e.g. chronic obstructive pulmonary disease, COPD)^1^ and autoimmune dysfunction,^2^ while effective removal of apoptotic cells helps drive resolution of inflammation.^3^ Given that macrophage interactions with apoptotic cells can be found in multiple human pathologies, we wished to understand how such interactions affect macrophage behaviour at a cellular level. Therefore, to study the effects of developmental and pathological levels of apoptosis on macrophages, we used *Drosophila* embryos, exploiting the ease of manipulating cell death and apoptotic cell clearance during *in vivo* imaging in this model.

*Drosophila* embryos contain a population of highly-motile macrophage-like cells (plasmatocytes, one of the three hemocyte cell types), that disperse to cover the entire embryo during development, removing apoptotic cells and secreting extracellular matrix as they migrate.^4^ In addition to their functional similarities with vertebrate white blood cells, fly blood cells share genetic specification via the action of GATA and Runx family members.^5^ Dispersal of embryonic macrophages is controlled through expression of PDGF/VEGF-related ligands (Pvfs) along their route,^6^^−^^8^ coupled with physical constraints.^9^ Cell-cell repulsion^10^ and downregulation of Pvfs^8^ contribute to the timing and stereotyped nature of later migratory events - lateral migration of macrophages from the ventral midline to the edges of the developing ventral nerve cord (VNC).

*Drosophila* macrophages prioritise their activites in the developing embryo with apoptotic cell clearance taking precedence over their dispersal.^11^ Surprisingly, prior exposure to apoptotic cells seems required for normal responses to injury and infection.^12^ Inflammatory responses to sterile injuries strongly resemble those observed in vertebrates with a Duox (also known as Cy) dependent burst in hydrogen peroxide essential for normal recruitment to sites of damage^13,14^ and a genetic requirement for specific Src family kinases within innate immune cells.^15,16^

During embryonic development, *Drosophila* macrophages work in concert with the glia of the central nervous system (CNS) to remove apoptotic cells.^17^ Both phagocytes utilise a variety of receptors to recognise and engulf apoptotic cells,^18^ one of which is Simu (also known as NimC4). Simu is a member of the Nimrod family of cell surface receptors with homology to members of the CED-1 family of receptors e.g. CED-1 (*C. elegans*),^19^ Draper (*Drosophila*),^20^ Jedi-1 and MEGF10 (mouse).^21^ Simu binds phosphatidylserine (PS) via an EMILIN-like domain at the N-terminus^22^ and its absence from macrophages and glia leads to a failure in removing developmentally-programmed apoptosis.^23^ In some contexts Simu may operate upstream of Draper,^23^ though direct physical interaction has not been demonstrated.

Here we used *simu* mutant fly embryos to challenge macrophages with pathological levels of apoptotic cell death to address how this affected their subsequent behaviour. We show that apoptotic cell death contributes to developmental dispersal of macrophages and that excessive amounts of apoptosis also induce defects in dispersal and migration. Chronic levels of uncleared apoptotic cells and acute induction of apoptotic cell death impair wound responses in vivo, importantly without a requirement for phagocytosis of these dying cells. Finally, in our new paradigm, we demonstrate a novel role for Simu in retention of macrophages at sites of necrotic wounds and reveal that Simu facilitates clearance of non-apoptotic cells at such sites of injury. Therefore, the role of Simu is not limited to clearance of apopotic cells, but regulates responses to damaged self in general.

## Results

### Apoptotic cells regulate developmental dispersal of macrophages

To understand the role of developmentally-programmed apoptosis in dispersal of macrophages, *Df(3L)H99* mutant embryos, which lack all apoptosis owing to deletion of the pro-apoptotic genes *hid*, *reaper* and *grim*,^24^ were examined. As per previous reports,^6,25^ macrophage dispersal was grossly normal in the absence of apoptosis (Fig. 1a-b). However, detailed analysis at stage 13 of development revealed that more macrophages were present in lateral positions in embryos that lacked apoptosis compared to controls (Fig. 1c-d). By stage 15, macrophage localisation on the VNC was similar in controls and embryos lacking apoptosis (Fig. 1e-f). Macrophage dispersal and VNC development are interdependent,^9^ but in this instance we saw no obvious VNC defects in the absence of apoptotic cell death,^26^ making this dispersal phenotype unlikely to be a consequence of morphogenetic defects (Fig. 1e-f). Consistently, live imaging of GFP-labelled macrophages at stage 13 revealed that significantly greater numbers of macrophages left the ventral midline and migrated laterally in the absence of apoptosis (Fig. 1g). Furthermore, this precocious migration was reflected in an enhancement in macrophage speed in the absence of apoptosis at stage 13, while speeds were similar to controls at stage 15 (Fig. 1h).

**Figure 1.**
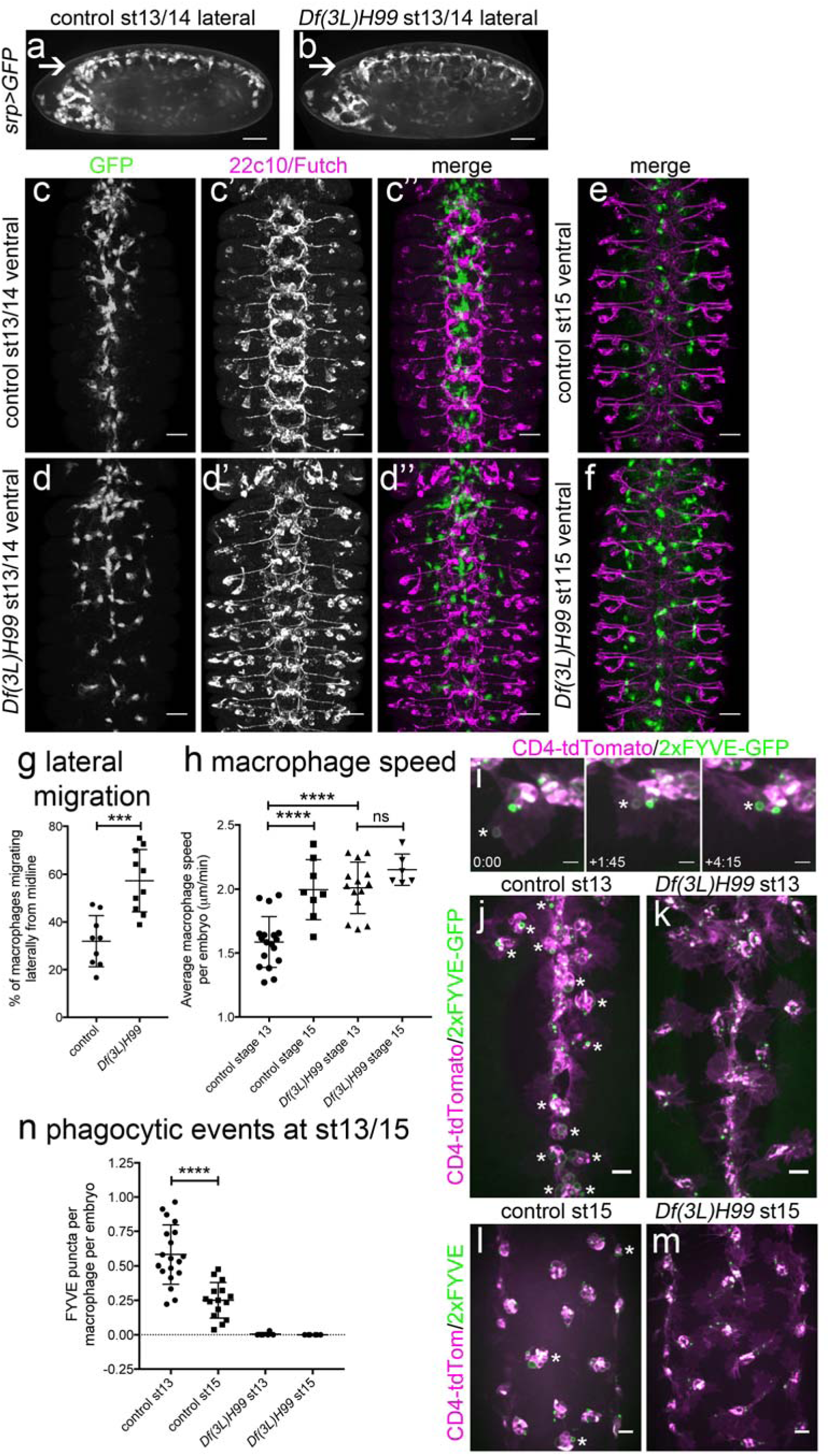
Apoptotic cell death contributes to developmental dispersal of *Drosophila* embryonic macrophages. (**a-b**) lateral projections (anterior is left) of control (*w;srp-GAL4,UAS-GFP*) and apoptosis-null *Df(3L)H99* mutant embryos (*w;srp-GAL4,UAS-GFP;Df(3L)H99*) showing macrophage distribution at stage 13/14 of development, including along ventral midline (arrow). (**c-d**) ventral projections (anterior is up) showing macrophages (anti-GFP staining, green in merge) and structure of the CNS (22c10/anti-Futch staining, purple in merge) at stage 13 of development in controls (*w;;crq-GAL4,UAS-GFP*) and in the absence of apoptosis (*w;;Df(3L)H99,crq-GAL4,UAS-GFP*). Macrophage projections constructed from z-slices corresponding to superficial macrophages on the ventral side of the VNC only, whereas Futch projection covers the entire volume of the VNC. (**e-f**) ventral projections (anterior is up) showing macrophage distribution (GFP) and CNS structure (Futch) at stage 15 in control and *Df(3L)H99* mutant embryos (genotypes as per **c-d**). (**g**) scatterplot showing percentage of macrophages on the midline that move laterally at stage 13 of development in control and apoptosis-null embryos (n=9 and 10, respectively; p=0.0006, Mann-Whitney test). (**h**) scatterplot showing speed per macrophage, per embryo (μm per min) in control and apoptosis-null embryos at stage 13 (p<0.0001) and stage 15 (n=18, 8, 14, 6 (left-right); p=0.47, one-way ANOVA with Tukey’s multiple comparison test). Genotypes in (**g-h**) are as per (**c-f**). (**i**) stills from movie of macrophages phagocytosing at stage 13 on the ventral midline in a control embryo (*w;srp-GAL4,UAS-2xFYVE-GFP;crq-GAL4,UAS-CD4-tdTomato*). EGFP and tdTomato are shown in green and purple, respectively, while asterisk shows nascent phagosome at indicated times (mins:secs) as it become positive for the PI3P sensor 2xFYVE-GFP post-engulfment; CD4-tdTomato labels membranes. (**j-m**) ventral projections of control (*w;srp-GAL4,UAS-2xFYVE-GFP;crq-GAL4,UAS-tdTomato*) and apoptosis-null embryos (*w;srp-GAL4,UAS-2xFYVE-GFP;crq-GAL4,UAS-tdTomato;Df(3L)H99*) at stage 13 (**j-k**) and stage 15 (**l-m**), showing recent phagocytic events via 2xFYVE-GFP sensor for PI3P. Asterisks show examples of 2xFYVE-GFP positive cells (**j**, **l**). (**n**) scatterplot showing number of phagocytic events (via 2xFYVE-GFP positive vacuoles) per macrophage, per embryo at stages 13 and 15 (n=19 and 15, respectively; p<0.0001, Mann-Whitney test). Central lines show mean and error bars represent standard deviation (**g-h,l**); *** and **** denote p<0.001 and p<0.0001; scale bars indicate 50μm (**a-b**), 10μm (**c-f**), 5μm (**i**) and 10μm (**j-m**).

By stage 15 of development the majority of apoptosis occurs within the CNS,^26^ and the formation of septate junctions by surface glia prevents macrophages from accessing these dying cells.^27^ Therefore, we predicted that macrophages at stage 15 would have fewer apoptotic cell phagosomes in comparison to stage 13. To test this, a phosphatidylinositol 3- phosphate (PI3P) sensor was expressed in macrophages (2xFYVE-GFP)^28^ to mark nascent phagosomes^29^ (Fig. 1i). Consistent with sequestration of apoptosis within the CNS, there were significantly fewer nascent phagosomes visible within macrophages at stage 15 compared to stage 13 (Fig. 1j, l, n), and such phagosomes were absent in apoptosis-null embryos (Fig. 1k, m-n). Therefore, it is only once macrophages become less exposed to apoptotic cell death that they become more motile, suggesting interactions with apoptotic cells restrict macrophages from migrating laterally. Taken together, these data suggest that interactions between developmentally-programmed apoptotic cell death and macrophages can delay lateral migration and contribute to the regulation of macrophage dispersal in *Drosophila* embryos.

### Pathological levels of apoptosis are associated with developmental defects in macrophage dispersal

Having identified a novel role for apoptotic cell death in regulating dispersal of *Drosophila* embryonic macrophages, we wished to investigate the consequences of pathological levels of apoptosis on macrophage behaviour *in vivo*. Embryonic macrophages migrate along the ventral midline in a constrained channel,^9^ in close contact with the epithelium and developing CNS (Fig. 2a-b). Later in development the environment becomes less constricted and macrophages disperse over the ventral side of the embryo, albeit still sandwiched between the epithelium and CNS (Fig. 2c-d). Thus, macrophages migrate in very close contact with the other main phagocyte population in the developing embryo: the glia of the VNC.

**Figure 2.**
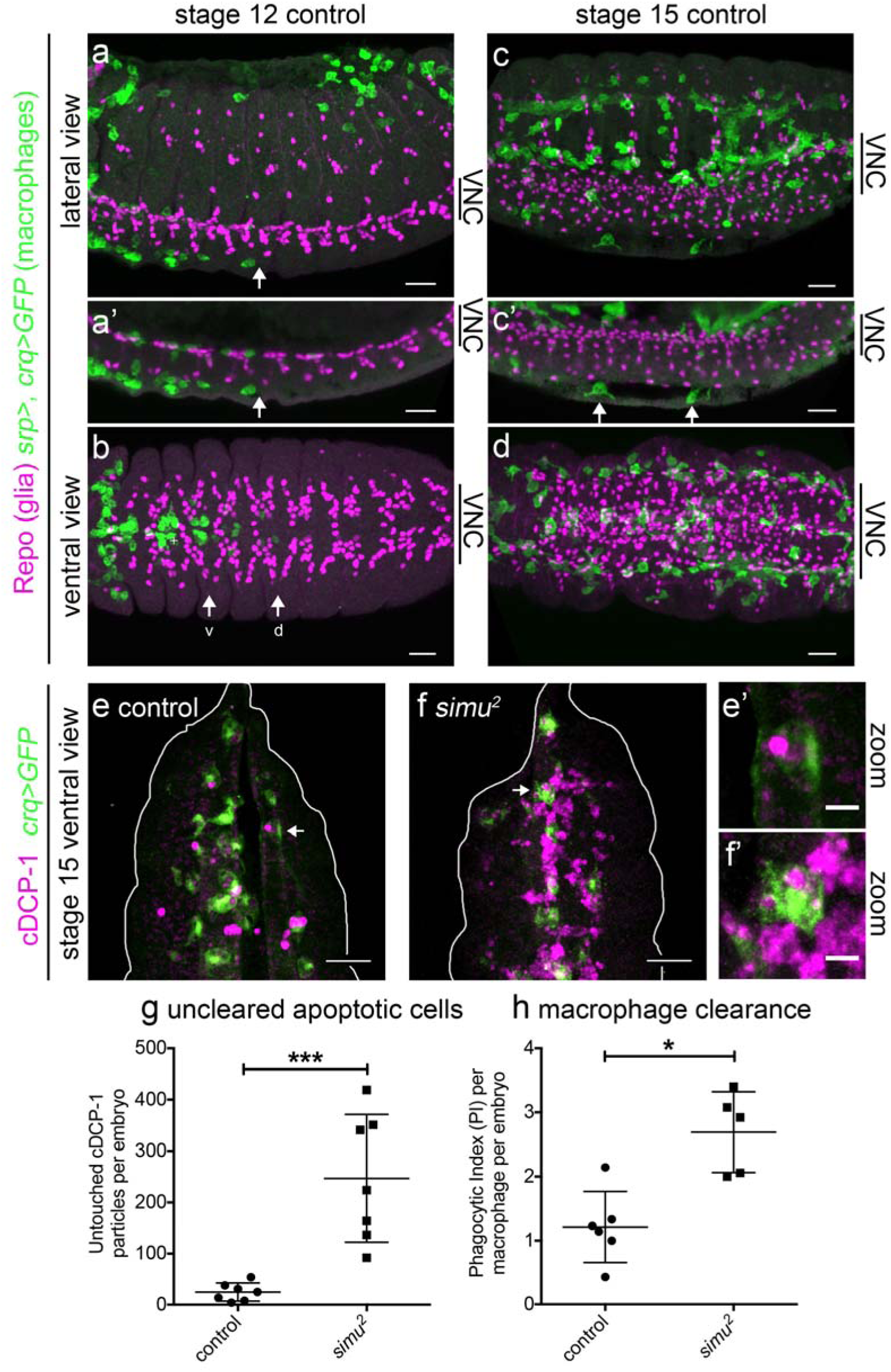
Loss of *simu* function results in exposure of macrophages to pathological levels of uncleared apoptotic cells. (**a-d**) maximum projections of control embryos (*w;srp-GAL4,UAS-GFP/+;crq-GAL4,UAS-GFP/+*) immunostained to show close contact of macrophages (anti-GFP, green), and glia (anti-Repo, purple) at stage 12 during migration along both sides of the ventral midline (**a**, lateral; **b**, ventral) and later in development at stage 15 (**c**, lateral; **d**, ventral). (**a’**) and (**c’**) are maximum projections of a smaller number of z-slices to show position of macrophages between epidermis and and VNC. Anterior is left and arrows show pioneer macrophages; arrows ‘v’ and ‘d’ in (**b**) label most advanced macrophages moving along the ventral and dorsal side of the ventral nerve cord (VNC), respectively; arrows in (**d’**) show two macrophages between VNC and epidermis. (**e-f**) maximum projections of stage 15 control and *simu* mutant embryos immunostained for GFP (green) and cleaved DCP-1 (cDCP-1, purple) to show macrophages and apoptotic particles, respectively. Projections are ventral views and correspond to void between epidermis and VNC. (**e’-f’**) show zooms of macrophages indicated by arrows in (**e-f**); white line shows edge of embryos. (**g**) scatterplot of untouched apoptotic punctae (cDCP-1 punctae not in contact with macrophages in the field of view) per embryo in control and *simu* mutant embryos at stage 15 (n= 7 per genotype; p=0.0006, Mann-Whitney test). (**h**) scatterplot of phagocytic index (cDCP-1 punctae engulfed per macrophage, per embryo) in control and *simu* mutant embryos at stage 15 (n=6 and 5, respectively; p=0.017, Mann-Whitney test). Genotypes in (**e-h**) are *w;;crq-GAL4,UAS-GFP* (control) and *w;simu^*2*^;crq-GAL4,UAS-GFP* (*simu* mutants). Error bars represent mean ± standard deviation; * and *** denote p<0.05 and p<0.001; scale bars indicate 10μm (**a-d**), 20μm (**e-f**) and 5μm in zoomed panels (**e’-f’**).

We predicted that removing the apoptotic cell clearance receptor Simu from macrophages and glia would lead to an overstimulation of macrophages with apoptotic cells, owing to a reduction in apoptotic cell clearance by these neighbouring cell populations. To confirm this prediction, apoptotic cell death was visualised by staining embryos using an antibody to cleaved DCP-1 (cDCP-1), an effector caspase, itself cleaved during apoptosis.^30^ Consistent with the role of Simu in apoptotic cell clearance, the absence of *simu* led to large numbers of apoptotic cells remaining unengulfed at stage 15 in the space between the epithelium and developing VNC, despite the proximity of macrophages, in contrast to controls, in which very few uncleared apoptotic cells persisted (Fig. 2e-f). Significantly, *simu* mutants exhibited a large increase in the number of untouched apoptotic cells in this region of the developing embryo (Fig. 2g), with only a small increase in the phagocytic index of macrophages (Fig. 2h). Untouched apoptotic cell punctae were used as a more conservative estimate of uncleared apoptotic cells, since macrophages were so overwhelmed that it was difficult to discern accurately whether phagocytosis had taken place from images of immunostained embryos. Therefore, in line with previous data,^23^ we could demonstrate a role for *simu* in apoptotic cell clearance, with the absence of *simu* function leading to pathological levels of apoptosis surrounding macrophages on the ventral side of the developing *Drosophila* embryo.

After establishing that loss of *simu* function leads to pathological levels of apoptosis in vivo, we sought to test the effect this exerted on macrophage behaviour. Migration of macrophages over the embryo is critical for embryonic development, since removal of dying cells and secretion of matrix are important aspects of morphogenesis.^31,32^ *Drosophila* macrophages clear apoptotic cells as they disperse throughout the embryo, contemporaneously with induction of apoptosis.^33^ To assess developmental dispersal we scored macrophage progression along the midline at stage 13 and numbers of macrophages on the ventral midline at stage 15 in control and *simu* mutant embryos. No gross defects were observed in progression of macrophages along the ventral midline at stage 13 in *simu* mutants compared to controls (Fig. 3a-c), nor was there a difference in the numbers of macrophages on the midline at stage 13 (Fig. 3d-f). However, by stage 15 there was a significant reduction in macrophage numbers on the ventral midline in *simu* mutants compared to controls (Fig. 3g-i). This suggested that the presence of excessive numbers of apoptotic cells can perturb dispersal of macrophages, consistent with the sensitivity of these cells to developmentally-programmed apoptosis (Fig. 1).

**Figure 3.**
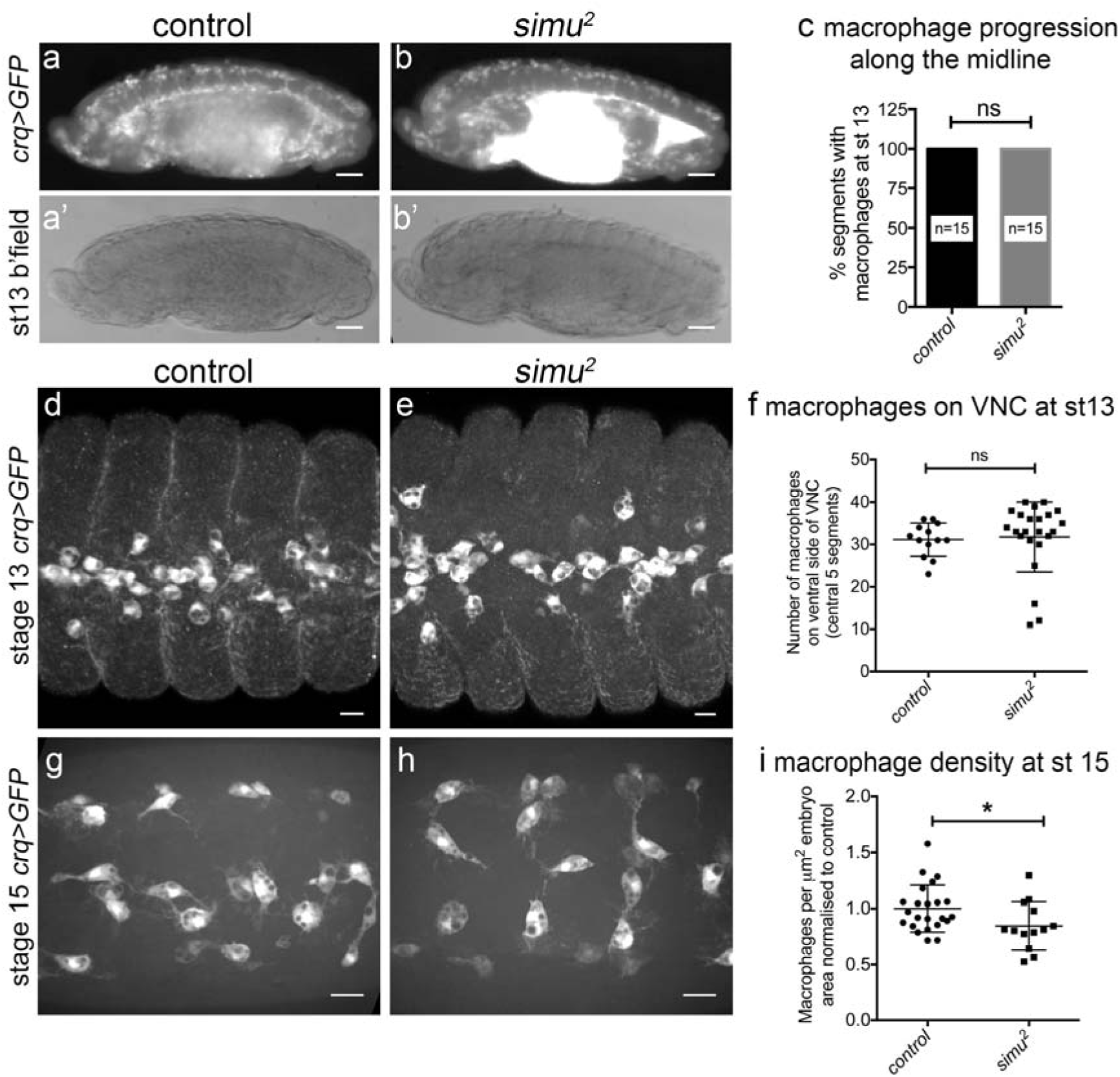
Loss of *simu* function leads to defects in developmental dispersal of macrophages. (**a-b**) lateral images of control (*w;;crq-GAL4,UAS-GFP*)and *simu* mutant embryos (*w;simu^*2*^;crq-GAL4,UAS-GFP*) showing migration along the ventral midline at stage 13; upper images show GFP channel (**a-b**), lower images are brightfield (**a’-b’**). (**c**) bar graph showing percentage of segments with macrophages on the ventral side of the VNC at stage 13 in controls and *simu* mutants (n=15 and 15, respectively; p>0.999, Mann-Whitney test). (**d-e**) ventral views of macrophages on the midline at stage 13 in control and *simu* mutant embryos immunostained for GFP. (**f**) scatterplot of numbers of macrophages on the ventral midline (central 5 segments) at stage 13, scored from anti-GFP immunostained embryos (n=13 and 23, respectively; p=0.15, Mann-Whitney test). (**g-h**) ventral views of GFP-labelled macrophages imaged live on the midline at stage 15 in control and *simu* embryos. (**i**) scatterplot of numbers of macrophages on the ventral midline at stage 15 (normalised according embryonic area in the field of view) in control and *simu* mutant embryos (n=23 and 13, respectively; p=0.040, Mann-Whitney test). All genotypes are as per (**a-b**). Error bars on graphs show standard deviation, line or bars show mean (**c**, **f**, **i**); ns and * denote not significant and p<0.05; scale bars indicate 50μm (**a-b**) and 10μm (**d-e**, **g-h**).

### Pathological levels of apoptosis impair basal migration of macrophages

Given the decreased numbers of macrophages on the ventral midline in the presence of excessive amounts of apoptosis (Fig. 3g-i), we wished to address whether the migration of these cells was perturbed in *simu* mutants. We measured basal cell motility by imaging the wandering migration of GFP-labelled macrophages at stage 15, finding that migration speeds were significantly reduced in *simu* mutants compared to controls (Fig. 4a-c; Supplementary movie 1). Despite the large numbers of uncleared apoptotic cells surrounding macrophages in *simu* mutants (Fig. 2e-g), there was only a minor stimulation of phagocytosis in *simu* mutants (Supplementary Fig. 1a-c). We were unable to detect obvious or repeated attempts to phagocytose apoptotic cells, suggesting that the defects in migration may not simply be a consequence of frustrated phagocytic events. Furthermore, macrophages exhibited a similar morphology to controls in *simu* mutants, exhibiting large, well-spread lamellipodia, with the only morphological difference detected being decreased circularity (Supplementary Fig. 1a-b, d-e). Therefore the motility machinery of these embryonic macrophages remains intact, consistent with a relatively mild defect in their dispersal (Fig. 3). As both basal migration and developmental dispersal were impaired in *simu* mutants, we next turned to an assay of inflammatory migration to address whether other pathologically-relevant macrophage behaviours were perturbed in the face of large amounts of uncleared apoptotic cells.

**Figure 4.**
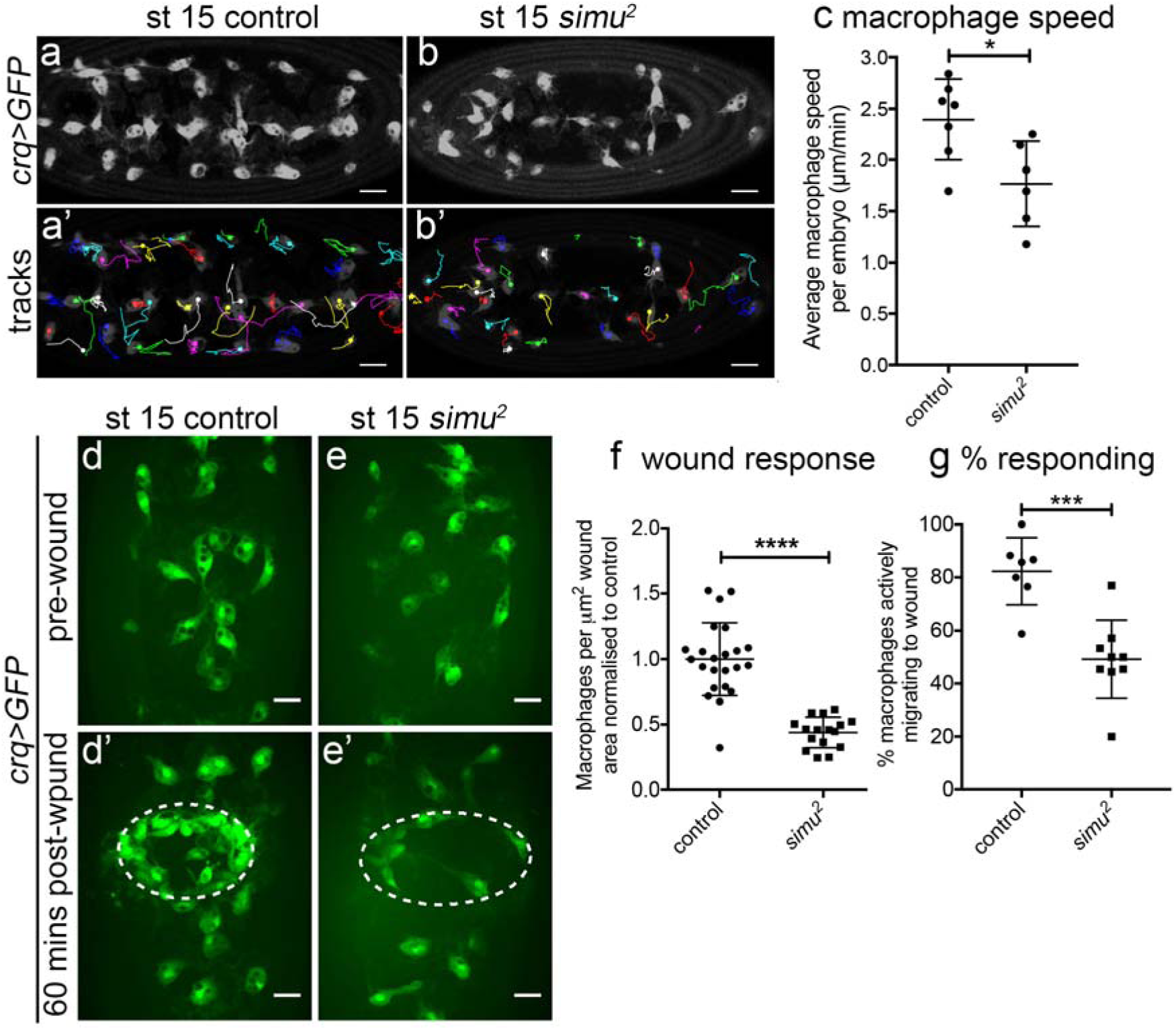
Pathological levels of apoptosis impair motility of macrophages and inflammatory migration to wounds. (**a-b**) images of GFP-labelled macrophages and their associated tracks on the ventral midline at stage 15 in control (*w;;crq-GAL4,UAS-GFP*, **a**) and *simu* mutant (*w;simu^*2*^;crq-GAL4,UAS-GFP*, **b**) embryos. (**a’-b’**) tracks show macrophage migration over a 30-minute period; dots show starting positions of those macrophages present in the first frame of the movie; anterior is left. (**c**) scatterplot of average speed per macrophage, per embryo on the ventral midline at stage 15 in μm per min in controls and *simu* mutants (n=7 and 6, respectively; p=0.035, Mann-Whitney test). (**d-e**) images of GFP-labelled macrophages on the ventral midline in stage 15 control (**d-d’**) and *simu* mutant (**e-e’**) embryos before (**d-e**) and at 60-minutes post-wounding (**d’-e’**). Dotted lines indicate wound sites. (**f**) scatterplot of wound response at 60 minutes (number of macrophages at wound divided by wound area, normalised to control average) in control and *simu* mutant embryos (n=23 and 16, respectively; p<0.0001, Mann-Whitney test). (**g**) scatterplot showing percentage of macrophages responding to wounds (percentage of those in field of view at t=0 not initially in contact with the wound site that reach the wound site in a 60-minute movie) in control and *simu* mutant embryos (n=7 and 9, respectively; p=0.0006, Mann-Whitney test). All genotypes as per (**a-b**). Lines represent mean, error bars represent standard deviation; *, *** and **** denote p<0.05, p<0.001 and p<0.0001 (**c**, **f**, **g**); scale bars indicate 20μm (**a-b**) and 10μm (**d-e**).

### Pathological levels of apoptosis impair inflammatory migration to wounds

Macrophages undergo a rapid and highly-directional inflammatory migration to laser-induced wounds in *Drosophila* embryos.^34^ Laser wounds induce calcium waves in the epithelium, leading to production of hydrogen peroxide, which is essential for efficient recruitment of macrophages to these sites of damage.^14^ Imaging GFP-labelled macrophages in *simu* mutant embryos revealed a significant defect in recruitment of macrophages to wounds (Fig. 4d-f; Supplementary movie 2). This effect was far greater than the small reduction in macrophage numbers present on the midline ahead of wounding in *simu* mutants (Fig. 3g-i), suggesting the defect in inflammatory migration is not purely due to a reduction in cells available locally to respond. Furthermore, the proportion of macrophages responding to wounds was also quantified, revealing that a significantly greater percentage of macrophages migrated to wounds in controls compared to *simu* mutants (Fig. 4g). Therefore, the ability to respond to wounds is compromised in *simu* mutants at a cellular level, potentially due to the presence of large numbers of uncleared apoptotic cells in the embryonic milieu. Imaging calcium responses following wounding, using the cytoplasmic calcium sensor GCamp6M,^35^ indicated that this reduction in inflammatory recruitment was not due to defects in the generation of wound cues (Supplementary Fig. 1f-i).

Wound recruitment defects are specifically associated with loss of *simu* function, since placing the loss-of-function allele *simu*^*2*^ in trans to a deficiency that deletes *simu* also perturbed inflammatory responses (Supplementary Fig. 1j-k). Furthermore, re-expression of wild-type *simu* in macrophages partially ameliorated *simu* mutant defects (Supplementary Fig. 1l-m), in line with a role for *simu* in both macrophage and glial-mediated apoptotic cell clearance.^23^

### Acute induction of apoptosis is sufficient to impair inflammatory responses

Since pathological levels of apoptosis appeared to impair wound responses in *simu* mutants, we wished to understand whether chronic or acute exposure to dying cells underlied this phenotype. Furthermore, carefully controlled introduction of apoptotic cell death would enable an understanding of whether efferocytosis was necessary for apoptosis-induced impairment of wound responses. To achieve this the pro-apoptotic regulator *hid* was overexpressed through a short heat-shock treatment of developing embryos that contained this gene under the control of the *hsp70* promoter (*hs-hid*)^36^ The length of heat-shock treatment was determined by imaging embryos containing ubiquitous expression of a caspase activity reporter (apoliner)^37^ and the *hs-hid* transgene (Fig. 5a and Supplementary Fig. 2a-d). Induction of apoptosis was also confirmed by staining for activated caspases (Fig. 5b).

**Figure 5.**
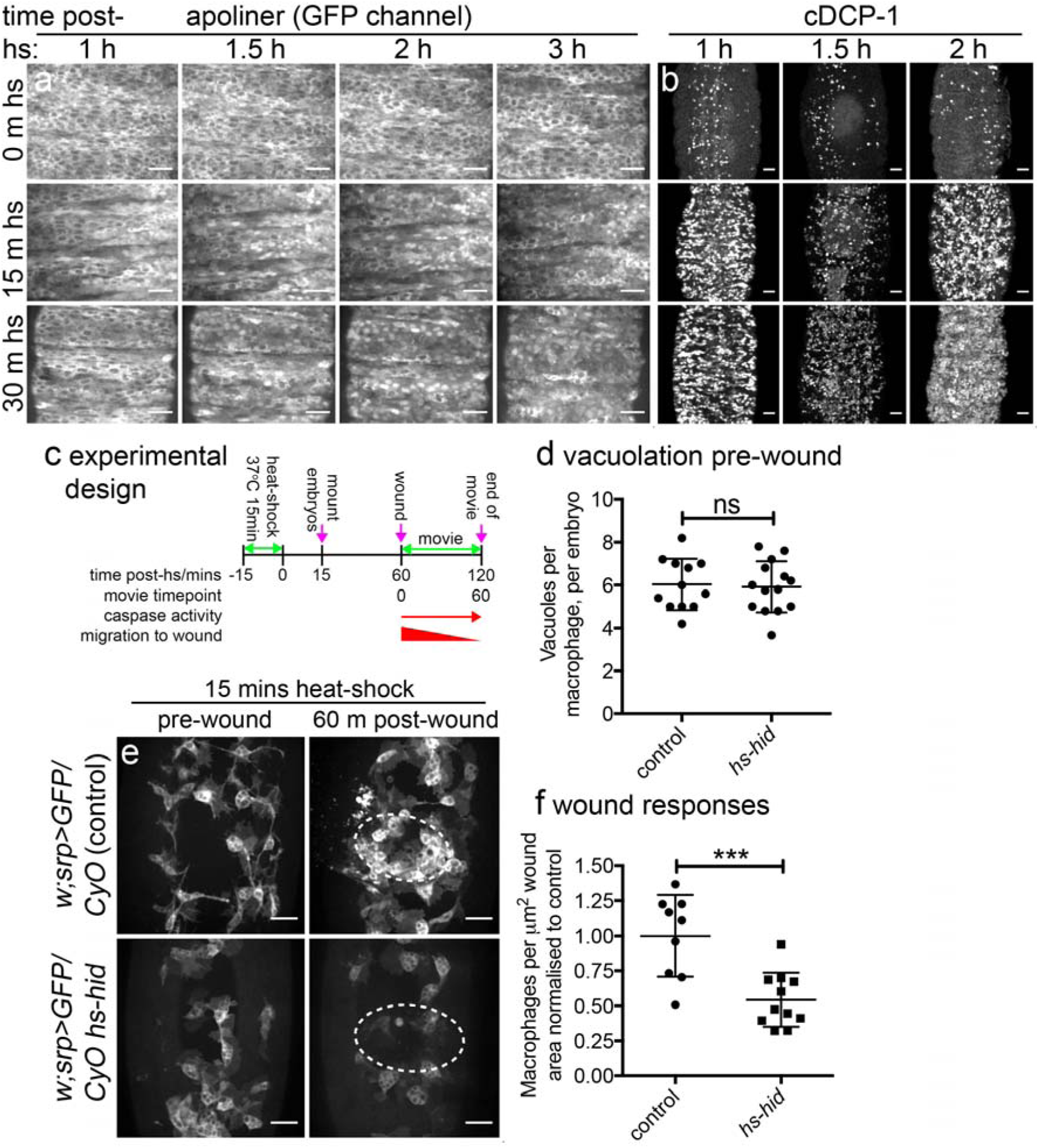
Acute induction of apoptosis impairs inflammatory responses without the requirement for engulfment by macrophages. (**a**) embryos containing ubiquitous expression of a caspase activity reporter (*w;CyO hs-hid/+;da-GAL4,UAS-apoliner/+*) heat-shocked for 0, 15 or 30 minutes and imaged from 60- minutes post-heat shock. Images show ventral projections of GFP channel taken from movies with nuclear localisation of GFP denoting presence of active caspase in that cell. Caspase activity apparent from 60-minutes post-heat shock; see Supplementary Fig. 2 for GFP and RFP channels and zooms. (**b**) ventral projections of controls (*w;srp-GAL4,UAS-GFP/+*) and embryos containing *hs-hid* (*w;CyO hs-hid/+*) stained for active caspases (anti-cDCP-1) following heat-shock for 0, 15 or 30 minutes; embryos fixed and stained at indicated times. (**c**) schematic showing experimental design to induce exogenous apoptosis ahead of clearance by phagocytes in the embryo. (**d**) scatterplot showing vacuoles per macrophage, per embryo as a read out of phagocytosis at 60-minutes post heat-shock (immediately before wounding) in control and *hs-hid* embryos (n=12 and 14, respectively; p=0.829, Mann-Whitney test). (**e**) pre-wound and 60-minutes post-wound ventral views of control (*w;srp-GAL4,UAS-GFP/CyO*) and *hs-hid* embryos (*w;srp-GAL4,UAS-GFP/CyO hs-hid*) subjected to 15-minute heat-shock; experimental design was as per (**c**), with wounding taking place at 60-minutes post heat-shock. Dotted lines show wound edges. (**f**) scatterplot of wound responses (density of macrophages at wound sites normalised to wound area and to control average) at 60-minutes post-wounding of control and *hs-hid* embryos (n=9 and 11, respectively; p=0.0005, Mann-Whitney test). Line and error bars represent mean and standard deviation in all scatterplots; ns and *** denote not significant and p<0.001; scale bars indicate 20μm.

Stage 15 embryos were heat-shocked for 15 minutes and immediately mounted for wounding. Wounding was performed 60 minutes after heat-shock, a time-point when caspase activity could be seen in those cells destined to die by apoptosis, but ahead of their delamination and engulfment by macrophages (Supplementary Fig. 2a-d). The majority of cells that undergo apoptosis appear to be superficial epithelial cells and these typically remain in the epithelium over the course of our 60-minute wounding experiments (60-120 minutes post-heat shock), with a wave of delamination eliminating them from the epithelium by 4.5 hours (± 0.3h standard deviation, n=9 movies; Supplementary movie 3). Calibrating the assay in this fashion meant it was possible to discern whether engulfment was a prerequisite for antagonism of inflammatory responses by apoptotic cell death (Fig. 5c). The lack of engulfment at this time point was confirmed via quantification of vacuoles in macrophages, which are present in the same numbers in macrophages that have been heat-shocked either in the presence or absence of *hs-hid* (Fig. 5d).

Imaging inflammatory responses of macrophages following induction of exogenous apoptosis revealed a significant defect in macrophage recruitment to wounds at 60-minutes post-wounding compared to control embryos (Fig. 5e-f; Supplementary movie 4). Importantly, no macrophages were observed to undergo apoptosis in these assays (Supplementary movie 4), suggesting these cells are somewhat tolerant to *hid* expression. Wounding the same embryos in the absence of a heat-shock failed to reveal a defect in wound responses (Supplementary Fig. 2f-g), indicating impaired wound responses were specific to the induction of exogenous apoptosis and could not have been caused by differences in genetic background (e.g. insertion site of *hs-hid*). Taken together this suggested that acute induction of apoptosis can attenuate wound responses, phenocopying *simu* mutant embryos in which macrophages are overwhelmed by uncleared apoptotic cells. Furthermore, since we could not detect an increase in phagocytosis at the time-point of wounding, this suggested that phagocytosis of apoptotic cells is not necessary for these phenotypes.

### Removal of apoptosis from *simu* mutants restores normal migration and improves macrophage responses to wounds

In order to confirm that excessive levels of uncleared apoptotic cells in *simu* mutants are responsible for the reduction in cell motility and defects in inflammatory responses to damaged tissue, the ability of cells to undergo developmentally-programmed apoptosis was removed from this mutant background using the *Df(3L)H99* deficiency.^24^ Comparing apoptosis-null *simu* mutants (i.e. *simu^*2*^;Df(3L)H99* double mutants) with *simu* mutants (*simu*^*2*^ single mutants) revealed a rescue of macrophage migration speed (Fig. 6a-b), whereas there was no difference in migration speed between macrophages in control and apoptosis-null embryos (Fig. 6a-b; Supplementary movie 5). As expected no vacuoles were seen in macrophages within apoptosis-null embryos, confirming absence of apoptosis and efferocytosis (Fig. 6a). Thus, migration defects in *simu* mutants are specifically due to apoptosis and not a subtle morphogenetic or macrophage-specification defect.

**Figure 6.**
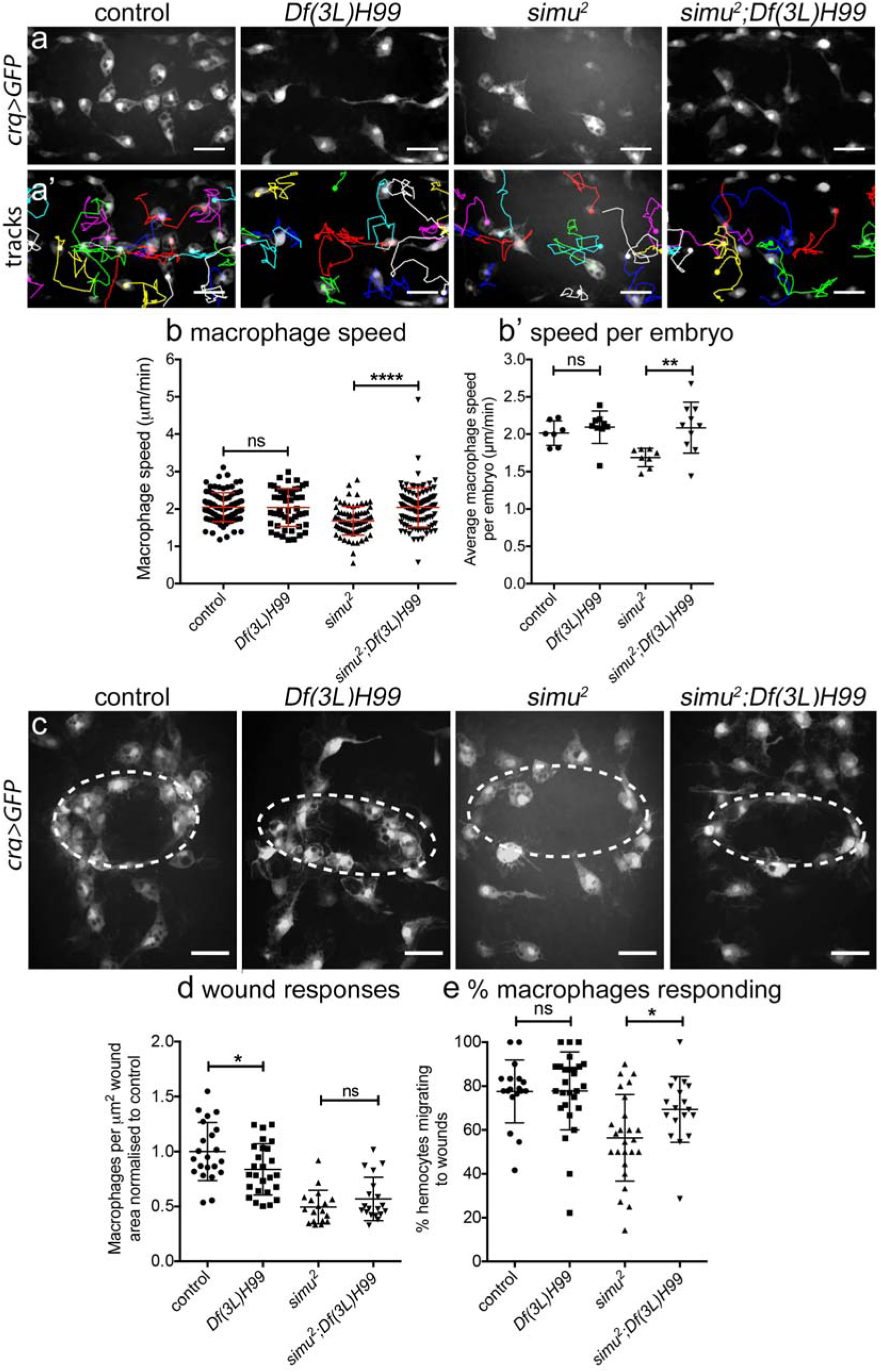
Removal of apoptosis from *simu* mutants rescues wandering migration and initial responses to wounds. (**a-a’**) maximum projections of GFP-labelled macrophages on the ventral midline at stage 15 (**a**) and tracks of their migration in the subsequent 60 minutes (**a’**) in controls, apoptosis-null embryos (*Df(3L)H99*), *simu* mutants (*simu*^*2*^) and *simu* mutants that lack apoptosis (*simu^*2*^;Df(3L)H99*). (**b-b’**) scatterplots of speed per macrophage (**b**) and average speed per embryo (**b’**) in μm per min at stage 15 in control and apoptosis-null embryos (n=7 and 9, respectively; p=0.252, Mann-Whitney test), and *simu* mutants and *simu* mutants that lack apoptosis (n=8 and 10, respectively; p<0.006, Mann-Whitney test, b’). (**c**) maximum projections of macrophages at wounds 60-minutes post-wounding in indicated embryos; dotted white ellipses show wound edges. (**d**) scatterplot of wound responses (macrophage density at wounds normalised to control average) comparing control and apoptosis-null embryos (n=22 and 26, respectively; p=0.039, Mann-Whitney test), and *simu* mutants and *simu* mutants that lack apoptosis (n=18 and 19, respectively; p=0.30, Mann-Whitney test). (**e**) scatterplot of percentage of responding macrophages (percentage that migrate to the wound that were not at the wound at t=0 minutes) comparing control and apoptosis-null embryos (n=18 and 27, respectively; p=0.80, Mann-Whitney test), and *simu* mutants and *simu* mutants that lack apoptosis (n=25 and 18, respectively; p=0.019, Mann-Whitney test). Genotypes for all embryos: control (*w;;crq-GAL4,UAS-GFP*), *Df(3L)H99* (*w;;Df(3L)H99,crq-GAL4,UAS-GFP*), *simu*^*2*^ (*w;simu^*2*^;crq-GAL4,UAS-GFP*) and *simu^*2*^;Df(3L)H99* (*w;simu^*2*^;Df(3L)H99,crq-GAL4,UAS-GFP*). Lines and error bars in scatterplots represent mean and standard deviation; ns, *, **, **** denote not significant, p<0.05, p<0.001 and p<0.0001; scale bars indicate 20μm.

Next we sought to test if removal of apoptosis from a *simu* mutant background might rescue inflammatory responses to sites of tissue damage. This represented a more difficult challenge since apoptosis-null embryos themselves exhibit a wound response defect,^12^ albeit presenting a less severe phenotype than *simu* mutants themselves (Fig. 6c-d). Quantifying the wound response at 60-minutes post-wounding showed no difference between *simu* and apoptosis-null *simu* mutant embryos (Fig. 6c-d). However, the percentage of macrophages responding to wounds showed a substantial rescue of this initial response to wounds in apoptosis-null *simu* mutants embryos compared to *simu* mutants (Fig. 6e; Supplementary movie 6). Thus, initial inflammatory responses to injury are normal in *simu* mutants in the absence of apoptosis, ruling out a specific role for *simu* in the detection or migration of macrophages to wounds. Instead, excessive apoptosis subverts normal macrophage responses to wounds at a distinct phase.

### Simu mediates phagocytosis of debris at wounds and is required to retain macrophages at sites of tissue injury

An abnormal number of macrophages were present at wounds 60 minutes after wounding in *simu* mutants devoid of apoptosis, despite a normal initial migratory response to wounds. Why then is there a defect in wound responses at later timepoints? Vertebrate innate immune cells are removed from sites of inflammation by either apoptosis^3,38^ or reverse migration,^39^^−^^42^ but there is no evidence to suggest that *Drosophila* macrophages die at wounds. Consequently, we investigated whether precocious exit from sites of damage explained the incomplete rescue of wound responses in *simu* mutants in the absence of apoptosis. Macrophages rarely left control wounds in the 60-minutes post-wounding; by contrast significantly more macrophages failed to remain at wounds in both *simu* and apoptosis-null *simu* mutant embryos compared to controls (Fig. 7a-b; Supplementary movie 7). Critically, this phenotype was maintained in the absence of apoptosis, suggesting that uncleared apoptotic cells were not drawing macrophages away from wounds, but rather *simu* regulates retention of macrophages at wounds. To our knowledge, this represents the first example of a gene controlling retention of macrophages at wounds in *Drosophila* embryos.

**Figure 7.**
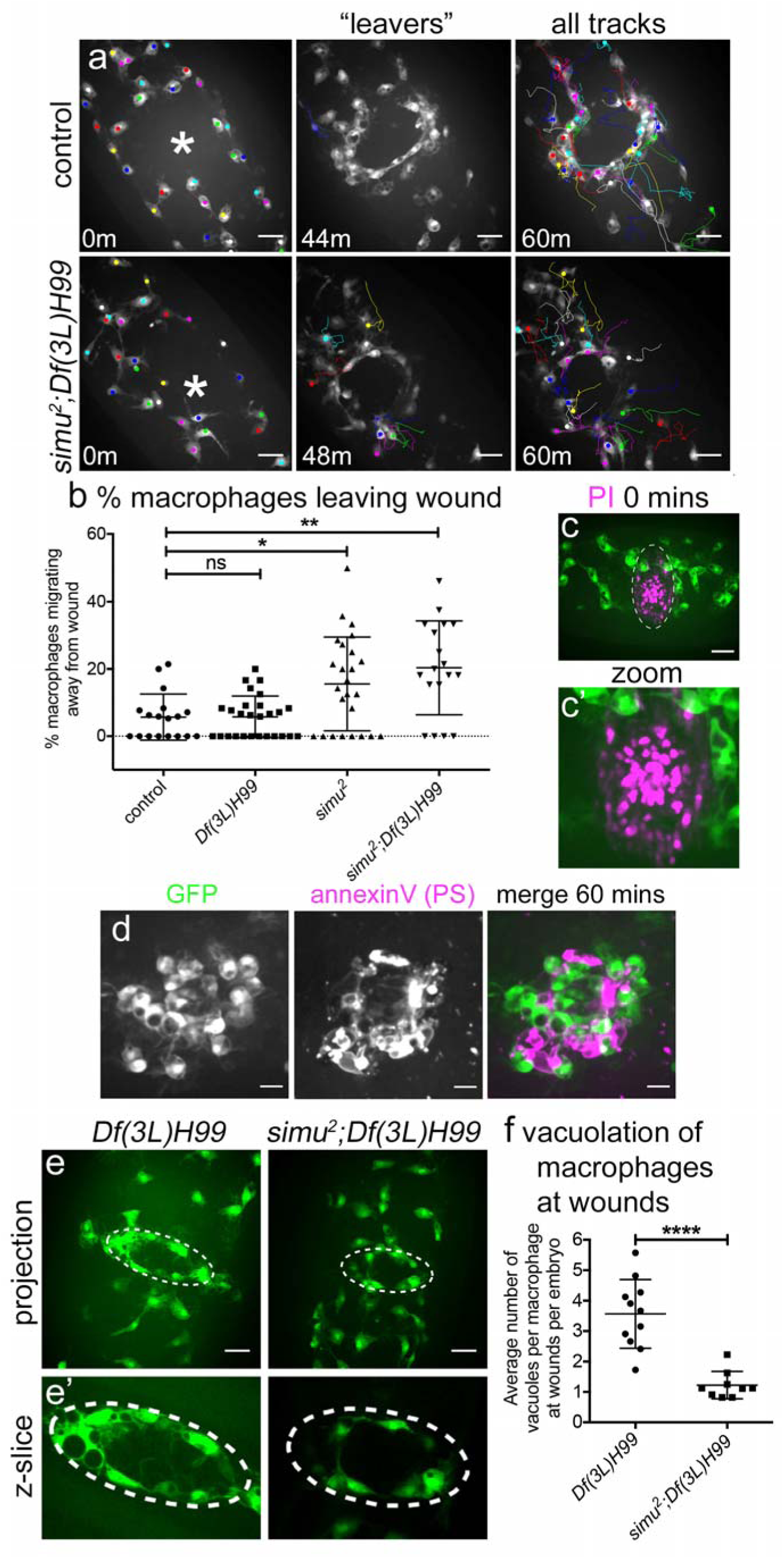
*simu* is required for phagocytosis of debris at wounds and to prevent precocious resolution of macrophages. (**a**) GFP-labelled macrophages and their associated tracks at 0, 44 or 48 and 60 minutes post-wounding in controls and *simu* mutants lacking apoptosis (*simu^*2*^;Df(3L)H99*). Left panels show macrophages present in field of view at 0 minutes; asterisk marks centre of wound. Central panels show tracks of macrophages that migrate to and then leave the wound (“leavers”). Right panels show tracks of cells present at 0 minutes that remain in the field of view until 60-minutes post-wounding. (**b**) scatterplot showing percentage of macrophages that migrate away from wounds over the course of 60-minute movies of inflammatory responses in controls, apoptosis-null embryos (*Df(3L)H99*), *simu* mutants (*simu*^*2*^), and *simu* mutants lacking apoptosis (*simu^*2*^;Df(3L)H99*); p values compared to compared to control are p=0.999, 0.047 and 0.031, respectively from left to right (Kruskal-Wallis test with Dunn’s multiple comparisons post-test; n=18, 27, 25, 18). (**c**) PI staining to show necrotic cells immediately after wounding in stage 15 control embryo (GFP-labelled macrophages, green; PI, purple). Dotted lines in upper panel show wound edge; lower panel shows zoom of wound. (**d**) panels show GFP-labelled macrophages, PS staining (via annexin V) and merged image at 60-minutes post-wounding of stage 15 control embryos. (**e**) maximum projections of GFP-labelled macrophages in apoptosis-null embryos (*Df(3L)H99*) and *simu* mutants that lack apoptosis (*simu^*2*^;Df(3L)H99*) at 60-minutes post-wounding; note absence of vacuoles in macrophages away from the wound (indicated via dotted line). (**e’**) shows zooms of wound regions taken from a single z-slices of z-stacks used to make projections in (**e**). (**f**) scatterplot showing numbers of vacuoles present in macrophages at wounds in apoptosis-null embryos (*Df(3L)H99*) and apoptosis-null embryos that also lack *simu* (*simu^*2*^;Df(3L)H99*) (n=11 and 9, respectively; p<0.0001, Mann-Whitney test). Genotypes as follows: control (*w;;crq-GAL4,UAS-GFP*), *Df(3L)H99* (*w;;Df(3L)H99,crq-GAL4,UAS-GFP*), *simu*^*2*^ (*w;simu^*2*^;crq-GAL4,UAS-GFP*), and *simu^*2*^;Df(3L)H99* (*w;simu^*2*^;Df(3L)H99,crq-GAL4,UAS-GFP*). Lines and error bars show mean and standard deviation in scatterplots; scale bars indicate 20μm (**a**, **c**, **e**) and 10μm (**d**) Images in (**c-d**) reproduced in Supplementary Fig. 3, which contains more detail for these experiments.

What then drives this Simu-dependent retention of macrophages? Laser ablation results in large amounts of necrotic death at wounds, as revealed by instantaneous entry of PI into ruptured cells at these sites of damage (Fig. 7c; Supplementary Fig. 3a-c). This loss of membrane integrity leads to externalisation of PS (Fig. 7d; Supplementary Fig. 3d), a known ligand of Simu.^22^ PS staining accumulates at sites of damage, particularly at the wound margin, and indicates the persistence of large amounts of debris, even at 60-minutes post-wounding (Fig. 7d; Supplementary Fig. 3d and movie 8).

We hypothesise that macrophages are retained at wounds through either inhibition of cell migration signals downstream of Simu or via physical interactions between PS and Simu at these sites of damage. Consistent with interaction of Simu and PS-positive necrotic debris at wounds, macrophages phagocytosed large amounts of debris at these sites in apoptosis-null embryos, but not in those embryos that lacked both *simu* function and apoptotic cell death (Fig. 7e-f). Consequently, these data reveal another novel function for Simu in that it is required for phagocytosis of non-apoptotic debris at wounds, suggesting that this receptor plays a more general role in clearance of damaged self in vivo, rather than acting specifically in efferocytosis.

## Discussion

Here we show that developmentally-programmed apoptosis contributes to dispersal of macrophages in fly embryos, while excessive amounts of apoptosis adversely impacts both macrophage migration and inflammatory responses to wounds. Acute induction of apoptosis phenocopied defects in inflammatory responses, suggesting apoptotic cells drive these changes in macrophage behaviour. Importantly, removal of apoptosis from a *simu* mutant background rescued migration and the initial inflammatory recruitment to wounds. This reveals a novel role for Simu in retention of macrophages at wounds and suggests that this receptor functions more generally to facilitate clearance of both apoptotic and necrotic cells.

We found that macrophage dispersal was impaired in the absence of apoptosis with precocious migration from the ventral midline, but also reduced numbers present on the ventral midline. Macrophage specification and migratory ability appeared normal in the absence of apoptosis, since cells were able to migrate at normal speeds. Dying cells may therefore provide instructive cues to aid dispersal. Indeed, apoptotic cells are known to play a role in recruitment of tissue-resident macrophages in other systems.^43,44^ Apoptotic midline cells^45^ may act to retain macrophages on the midline in wild-type embryos, hence explaining why macrophages migrated precociously from the midline in the absence of apoptosis. Alternatively, factors released by ‘undead’ cells (cells induced to die, but in which execution of apoptosis is blocked) may interfere with macrophage dispersal in the absence of apoptosis. Indeed, ‘undead’ cells are known to release signalling molecules such as TGFβ family proteins and H_2_O_2_,^46,47^ known regulators of macrophage behavior in many disease states that are conserved across different models.^14,48^ In contrast, uncleared apoptotic cells only exert later defects in dispersal, potentially since apoptotic cells must first accumulate to impact migration. Morphological defects are unlikely to contribute, since *simu* mutants appear grossly normal and are viable.^23^

We found that apoptotic cell death was necessary for impairment of migration and wound responses in *simu* mutants, with large amounts of uncleared apoptotic cells surrounding macrophages. Furthermore, we could reproduce wound recruitment defects by acute induction of apoptosis with macrophages failing to migrate to wounds. Phagocytosis of dying cells was not required for this effect, consequently we favour a model whereby ‘find-me’ signals released from uncleared apoptotic cells act to subvert normal macrophage responses. Find-me cues are signals released by dying cells to recruit phagocytes to facilitate clearance,^49^ though none have been identified to date in *Drosophila*. The release of find-me cues, or other stress signals produced in response to uncleared apoptotic cells, may confuse chemotactic responses of macrophages.

In the absence of wounds, we found that macrophages migrated more slowly in *simu* mutants, suggesting a general suppression of motility. This slowed migration seems unlikely to account for impaired wound responses, since macrophages lacking β-integrin move more slowly, but still ultimately reach wounds in normal numbers.^50^ In addition to the large increase in uncleared apoptotic cells, there was also a small increase in the phagocytic index of macrophages, although this only equated to approximately one extra puncta per cell. Others have reported that vacuolation of innate immune cells is associated with defective migration,^51,52^ while overloading macrophages with engulfed cargoes can lead to phagosomal maturation defects.^53^ Furthermore, other professional phagocytes such as *Dictyostelium* amoebae and dendritic cells pause migration when internalising cargo, albeit through macropinocytic mechanisms.^54,55^ However, we did not observe obvious repeated failures of phagocytic events by *simu* mutant macrophages. Macrophages in *simu* mutants are significantly less vacuolated than *SCAR* or *WAVE* complex mutant *Drosophila* macrophages and lysosomal storage mutant macrophages in zebrafish larvae.^51,52^ Whether subtle changes in the numbers of phagosomes seen in this study can account for such potent disruption of macrophage behaviour seems unlikely from the point of view of evolution of a functional innate immune system, but the threshold at which phagosomal maturation becomes pathological has yet to be directly addressed. An alternative explanation would be promotion of an anti-inflammatory state of programming of macrophages by apoptotic cell clearance.^56^ Currently, there is no strong evidence for such macrophage programming in *Drosophila*, though some progress has been made by Ando and colleagues.^57,58^ Nonetheless, clearance of apoptotic cells is associated with increased expression of phagocytic receptors in a number of phagocytic cell types, including macrophages.^12,59−61^ *Drosophila* embryonic macrophages also undergo remarkable shifts in behavior over the fly life cycle, controlled in part by lipid hormone signalling.^62,63^ However, the anti-inflammatory phenotype associated with acute induction of apoptosis would necessitate rapid reprogramming, unlikely for lipid hormone signalling, suggesting a more rapid mechanism, such as release of find-me cues.

Laser wounding of the epithelium led to accumulation of necrotic debris at sites of injury, including an accumulation and persistence of a marker used by phagocytes to clear apoptotic cells (PS). Simu possesses a PS-binding domain^22^ and is necessary for normal phagocytosis of necrotic debris at wounds. This work suggests that Simu is a receptor for both apoptotic and necrotic cells, and should be reclassified as a more general scavenger receptor, as for the *Drosophila* receptors Draper and Croquemort.^59,64,65^

Removal of uncleared apoptotic cells from a *simu* mutant background rescued migration and improved responses to wounds. The early departure of macrophages from wounds, which resembles reverse migration of neutrophils^39,42^ or exit of macrophages, which can leave sites of inflammation via lymphatic vessels in vertebrates,^41,66^ contributed to a partial rescue of wound responses. Simu may directly mediate retention of macrophages at wounds, potentially via binding to PS and/or other ligands, or it may act more indirectly possibly signalling through other co-receptors. The fact that initial responses to wounds were rescued in the absence of apoptosis implies that Simu is not interpreting chemotactic cues released from wounds, while precocious exit of macrophages from wounds in the absence of both Simu and apoptotic cell death (*simu^*2*^;Df(3L)H99* mutants) suggests these phagocytes are not simply being distracted away by apoptotic cells.

In summary, we show that the presence of apoptotic cells significantly alters macrophage behaviour in vivo and reveal new functions for the apoptotic cell receptor Simu. Our finding that pathological levels of apoptosis impair multiple macrophage functions has significant implications for a wide range of human conditions in which aberrant or subverted macrophage behaviour drives or exacerbates disease progression, including cancer, atherosclerosis, neurodegenerative disorders and other chronic inflammatory conditions.

## Materials and methods

### Fly genetics and husbandry

*Drosophila melanogaster* fruit flies were reared on standard cornmeal/agar/molasses media at 18°C or 25°C. Embryos were harvested from laying cages with apple juice agar plates left overnight at 22°C. *Srp-GAL4* and/or *crq-GAL4* were used to label macrophages with GFP, red stinger and/or CD4-tdTomato, or to drive expression of other transgenes. The following alleles, transgenes and deficiencies were used in this study: *srp-GAL4*,^7^ *crq-GAL4*,^34^ *da-GAL4*,^67^ *UAS-GFP*, *UAS-red stinger*,^68^ *UAS-CD4-tdTomato*,^69^ *UAS-2xFYVE-GFP*,^28^ *UAS-apoliner*,^37^ *UAS-GCamP6M*,^35^ *UAS-simu*,^23^ *simu*^*2*^,^23^ *Df(3L)H99*,^24^ *Df(2L)BSC253*,^70^ *CyO hs-hid*.^36^ Selection against the fluorescent balancers *CTG*, *CyO dfd*, *TTG* or *TM6b dfd* was used to discriminate homozygous mutant embryos for recessive lethal alleles.^71,72^ See Supplementary Table 1 for a full list of genotypes used in this study and their sources.

### Imaging and wounding of *Drosophila* embryos

Embryos were washed off apple juice agar plates, dechorionated in bleach, then washed in distilled water. Live and fixed embryos were mounted on slides in voltalef oil (VWR) or DABCO (Sigma) mountant, respectively, as per Evans et al., 2010.^9^ Immunostained embryos were imaged on a Nikon A1 confocal system using a 40X objective lens (CFI Super Plan Fluor ELWD 40x, NA 0.6), which was used for the migration movies in Fig. 4. All other time-lapse imaging and wounding was performed on a Perkin Elmer UltraView Spinning Disk system using a 40X objective lens (UplanSApo 40x oil, NA 1.3). Lower magnification images of embryos were taken using a MZ205 FA fluorescent dissection microscope with a PLANAPO 2X objective lens (Leica) or a 20x objective (UplanSApo 20x, NA 0.8) on the Perkin Elmer UltraView Spinning Disk system.

### Fixation and immunostaining of *Drosophila* embryos

Embryos from overnight plates were fixed and stained as per Evans et al., 2010.^9^ Antibodies were diluted 1:1 in glycerol for storage at −20°C and these glycerol stocks were diluted as indicated in PBS (Oxoid) containing 1% BSA (Sigma) and 0.1% Triton-X100 (Sigma). Rabbit anti-cleaved DCP-1 (1:500; 9578S – Cell Signaling Technologies), 22C10 mouse anti-Futch, (supernatant used at 1:100), 8D12 mouse anti-Repo, (concentrate used at 1:500 – Developmental Studies Hybridoma Bank) were used as primary antibodies at the indicated dilutions. Rabbit anti-GFP (1:500; ab290 – Abcam) or mouse anti-GFP (1:100; ab1218 – Abcam) were used to stain GFP-expressing macrophages for co-immunostaining with mouse and rabbit primary antibodies, respectively. Fluorescently-conjugated goat anti-mouse or goat anti-rabbit secondary antibodies (Alexa Fluor 568, Alexa Fluor 488 - Life Technologies, or FITC - Jackson Immunoresearch) were used to detect primary antibodies at 1:200.

### Macrophage migration assays

Embryos with fluorescently-labelled macrophages were mounted ventral-side up and left to acclimatise on slides for 30 minutes before imaging for all migration assays. For analysis of lateral migration (stage 13 embryos) and wandering/random migration (stage 15 embryos) z-stacks of macrophages between the epithelium and CNS were collected every 2 minutes for 1 hour (with the exception of data in Fig. 4 which was collected over a 30-minute period). Maximum projections of despeckled z-stacks were assembled in Fiji.^73,74^ Macrophages lying between the edges of the VNC at the start of the movie were tracked until they disappeared from view using the manual tracking plugin in Fiji.

For the wounding assay, pre-wound z-stacks were taken prior to wounding of stage 15 embryos on their epithelial surface using a nitrogen-pumped Micropoint ablation laser (Andor), as per Evans et al., 2015.^16^ Z-stacks of wounded embryos were collected every 2 minutes for 1 hour. At the end of the movie an additional z-stack was taken incorporating a brightfield image with which to measure the wound size. Movies were assembled as per wandering migration movies. The number of macrophages at wounds was quantified from a 15μm deep z-stack, with the first z-slice used in this analysis corresponding to the one containing the most superficial macrophage adjacent to, but not at the wound. Wound area was measured from the brightfield image taken at 60-minutes post-wounding. Macrophage responses to wounds were calculated as follows: number of macrophages within or touching the wound area divided by wound area in μm^2^. This was normalised according to control responses.

To calculate the initial inflammatory response to wounds the percentage of responding macrophages was calculated: this is the percentage of macrophages present at in the field of view at 0 minutes post-wounding that migrated to the wound; macrophages already present at the wound site at this time-point were not counted as responding, nor included in the total numbers present in the field of view. Macrophages that left the wound were not counted a second time if they returned and were defined as responding if they moved towards and touched the wound edge.

To assess resolution of macrophages from wounds, the percentage of macrophages that left the wound site was quantified. This is the proportion of macrophages (either that were present at the wound site at t=0 minutes or that reached the wound at any point during the wound movie) that left the wound during the course of a 60-minute time-lapse movie. N.B. if any part of a macrophage retained contact with the wound edge, it was not scored as having left.

### Introduction of exogenous apoptosis via heat-shock

*Hs-hid* or control embryos from overnight plates were washed off into embryo baskets (70μm cell strainers, Fisher) using distilled water. Baskets containing embryos to be heat-shocked were placed into a large weighing boat containing 75mL of pre-warmed distilled water in a circulating water bath set to 39°C for the indicated time. Non heat-shocked controls were incubated in distilled water at room temperature. Following heat-shock, embryos were dechorionated and then stage 15 embryos selected and mounted for live imaging 30 minutes before they were due to be wounded, such that they could acclimatise on slides. For instance, if embryos were wounded at 60-minutes post-heat shock, slides were set up to be ready for 30-minutes post-heat shock. Embryos were then wounded and imaged as described above to visualise macrophage behaviour or caspase activity via the apoliner reporter.^37^ Alternatively embryos were fixed at various time-points post-heat shock to reveal caspase activity via cDCP-1 immunostaining.

### Injection of propidium iodide and annexin V

Propidium iodide (PI) and annexin V are well-characterised reagents used to detect membrane permeabilisation/necrosis and exposure of PS/apoptosis, respectively.^75,76^ Embryos were mounted ventral-side up on double-sided sticky tape (Scotch), dehydrated for 7-10 minutes in an air-tight box containing sachets of silica gel (Sigma), then covered in voltalef oil before microinjection. Borosilicate glass needles (World Precision Instruments; capillaries were 1mm and 0.75mm inner and outer diameter capillaries, respectively) for microinjection were made using a P1000 needle puller (Sutter). Embryos were microinjected anteriorly into the head region with 1 mg/mL PI (Sigma) dissolved in PBS, or into the vitelline space with undiluted Alexa Fluor 568-labelled Annexin V (Invitrogen). After injection, a coverslip (thickness 1, VWR) was applied, supported by two coverslip bridges (thickness 1, VWR) placed on either side of the embryos, and fixed in place with nail varnish. Embryos were then imaged immediately, wounded, and then re-imaged to detect membrane permeabilisation and/or externalisation of PS. As a positive control for necrotic cell death, embryos were injected with PI, coverslips attached and then heated at 60°C in a humidified box for 15 minutes.

### Image processing, quantification and statistical analyses

Image processing was carried out in Fiji/ImageJ, with images despeckled in Fiji before maximum projections were assembled. Brightness and contrast enhancement was applied equally across all data sets being compared with each other. A Python (Python Software Foundation) script provided by the Whitworth Lab (University of Cambridge) was used to blind images ahead of analysis. Figures were assembled in Photoshop (Adobe).

Statistical analyses were performed using Prism (GraphPad); see legends and text for details of statistical tests used. N numbers refer to individual *Drosophila* embryos (unless otherwise stated), taken from laying cages containing >50 adult flies; experiments were conducted across at least three imaging sessions with control embryos mounted on the same slides each time. Previous experimental data shows that >6 movies (wandering migration) and 15-20 wound movies are sufficient to detect an effect size of 20% that of control values. Movies that lost focus or wounds that leaked obscuring macrophages at wounds were excluded from analyses. Immunostaining was performed on batches of pooled embryos collected across multiple days, with controls stained in parallel using staining solutions made up as a master mix and split between genotypes. All relevant raw data is available on request from the authors.

### Quantification of phagocytosis and failed apoptotic cell clearance

Anti-cDCP-1 and anti-GFP immunostained embryos were imaged on a Nikon A1 system using the 40X objective lens with a spacing of 0.25μm between z-slices. Channels were merged to enable discrimination of cDCP-1 punctae within/outside macrophages. A portion of the z-stack representing a 10μm deep volume corresponding to the area occupied by macrophages between the epithelium and VNC was used to count numbers of cDCP-1 punctae per macrophage to quantify phagocytic index. Only macrophages completely in view in the 10μm sub-stack were included in the analysis and these values were averaged to give an overall value per embryo. Untouched apoptotic cells were quantified by counting the number of red cDCP-1 punctae that are located within the 10μm sub-stack, but that were not in contact with GFP-labelled macrophages. This method was selected to enable comparison with the quantification of Kurant et al., 2008^23^ and also since it was technically challenging to discriminate whether cDCP-1 punctae in close contact with macrophages were inside or outside of macrophages in *simu* mutants owing to the extreme build up of uncleared apoptotic cells in this background.

Number of vacuoles per macrophage (averaged per embryo) was used as an indirect readout of phagocytosis. *Drosophila* macrophages exclude cytoplasmic GFP from phagosomes and these can be shown to correspond to apoptotic cells by their absence from macrophages in apoptosis-null *Df(3L)H99* embryos. Vacuoles were counted using z-stacks of GFP-labelled taken from live imaging experiments. Vacuoles were assessed in the z-slice in which each macrophage exhibited its maximal cross-sectional area. Therefore this analysis should not be considered as the absolute numbers of corpses per cell but a relative read-out thereof.

Phagocytosis was also quantified from stills or movies of macrophages expressing either 2xFYVE-GFP or CD4-tdTomato. Z-stacks were taken every 15 seconds for 20 minutes. Phagocytic events were scored when new phagocytic events occurred via formation of phagocytic cups (CD4-tdTomato) or the numbers of 2xFYVE-GFP punctae larger than 2μm with an obvious lumen present in macrophages, as a read-out of recently-formed phagosomes. 2xFYVE-GFP punctae of this size are absent from *Df(3L)H99* mutants.

### Analysis of developmental dispersal

Developmental dispersal of macrophages was analysed by scoring the presence of GFP-labelled macrophages between the epithelium and VNC in each segment at stage 13/14 following fixation and immunostaining for GFP. Embryos with macrophages in all segments were scored as 100% progression along the midline; an embryo that lacked macrophages in segments 6-9 would have been scored as 66% progression.

To quantify dispersal in more detail at stage 13, the number of macrophages present on the ventral side of the VNC (5 most-medial segments) was counted from images taken from a Nikon A1 confocal microscope. Numbers of macrophages on the ventral side of the VNC in pre-wound images was used to quantify macrophages numbers at stage 15; this was expressed as macrophage density, whereby number of macrophages was normalised according to the area in the field of view, as demarcated by the vitelline membrane from the least superficial (most dorsal) z-slice in the z-stack.

## Author Contributions

All authors contributed to planning of experiments and preparation of the manuscript. HGR, ELA and IRE performed and analysed the experiments. IRE conceived the project with intellectual input from SAJ.

## Acknowledgements

IRE is a Sir Henry Dale Fellow funded by Wellcome/Royal Society grant 102503/Z/13/Z. SAJ was supported by Medical Research Council and Department for International Development Career Development Award Fellowship (MR/J009156/1) and was additionally supported by a Krebs Institute Fellowship and Medical Research Council grant (G0700091).

This work would not be possible without the Bloomington *Drosophila* Stock Center (NIH P40OD018537) and Flybase (NIH and MRC grants U41 HG000739 and MR/N030117/1, respectively). Imaging was performed in the Wolfson Light Microscopy Facility using a Perkin Elmer spinning disk imaging system (MRC grant G0700091 and Wellcome grant 077544/Z/05/Z) and Nikon A1/TIRF system (Wellcome grant WT093134AIA). We thank Will Wood (University of Edinburgh), Martin Zeidler, Darren Robinson and Fly Facility staff (University of Sheffield) for their support and Steve Renshaw and Phil Elks (University of Sheffield) for critical reading and feedback on the manuscript.

